# Generative genomics accurately predicts future experimental results

**DOI:** 10.1101/2025.09.08.674753

**Authors:** Gregory Koytiger, Alice M. Walsh, Vaishali Marar, Kayla A. Johnson, Max Highsmith, Alexander R. Abbas, Andrew Stirn, Ariel R. Brumbaugh, Alex David, Darren Hui, Jeffrey M. Kahn, Sheng-Yong Niu, Liza J. Ray, Candace Savonen, Stein Setvik, Jeffrey T. Leek, Robert K. Bradley

## Abstract

Realizing AI’s promise to accelerate biomedical research requires AI models that are both accurate and sufficiently flexible to capture the diversity of real-life experiments. Here, we describe a generative genomics framework for AI-based experimental prediction that mirrors the process of designing and conducting an experiment in the lab or clinic. We created GEM-1 (Generate Expression Model-1), an AI system that effectively models the enormous range of bulk and single-cell gene expression experiments performed by scientists and benchmarked its performance across multiple biological axes. GEM-1’s prediction of future gene expression experiments–RNA-seq data deposited in public archives after our training data cutoff–yielded accuracy comparable to the best-possible performance estimated by comparing the results of matched lab experiments. Overall, our approach illustrates the transformative potential of generative genomics for applications ranging from predicting cellular perturbations in vitro to *de novo* generation of data from large clinical cohorts.

## INTRODUCTION

Although technological advances, such as highly multiplexed assays, continue to increase the power and speed of biological research, almost all lab-based experiments and clinical trials are limited by fundamental constraints. Many lab experiments can be conducted no faster than the speed with which cells grow or diseases develop, while clinical trials are similarly governed by the availability and recruitment of participants and essential regulations to protect patients.

The generative capabilities of modern artificial intelligence (AI) models with deep learning architectures offer the possibility of circumventing these fundamental biological constraints by computationally simulating many, or even all, limiting experimental steps. This vision is fast becoming reality for protein structures, where deep learning models excel at not just predicting the structure of existing proteins^1–4^ but also designing novel protein structures that do not occur in nature^5–8^. The protein structure problem has several features that have proven important for developing accurate AI models. First, protein structure prediction and design can be readily articulated as a well-defined problem with relatively simple inputs and outputs. Second, the Protein Data Bank provides an excellent source of curated data for training AI models. Third, the conservation of structure and function despite evolutionary sequence divergence across organisms that typifies most proteins means that genome sequencing across the evolutionary tree provides a vast source of information for training AI models^9–11^.

Like solving protein structures, measuring gene expression with RNA-seq is a foundational assay that is critical to modern biomedical research. However, several features of the gene expression problem render it less readily amenable to AI modeling. For example, a gene expression assay has a well-defined output (expression of all genes) but may not have an easily defined input (samples can come from virtually any conceivable lab experiment or clinical trial); public gene expression databases such as the National Center for Biotechnology Information Sequence Read Archive (NCBI SRA)^12^ contain vast gene expression archives but only inconsistently curated information on the experiments these data are derived from; and gene expression is less conserved across species than is protein structure^13,14^, so the natural laboratory of evolution provides limited information.

Despite these challenges, there is a recent explosion of interest in building deep learning models of gene expression. Most groups have focused on modeling single-cell RNA-seq (scRNA-seq) data, a natural choice given the availability of large atlases of curated scRNA-seq data that has been uniformly processed and harmonized^15,16^, facilitating the creation of well-defined inputs and outputs that are essential to training and testing a deep learning model.

These approaches are exciting and offer promising solutions to important scRNA-seq problems ranging from cell type annotation to perturbation prediction^17–23^. However, their focused architectures also constrain the variety of experiments that they can model.

Here, we undertook a different approach. We sought to construct an AI system that captures the vast diversity of gene expression experiments conducted in the biomedical enterprise, including basic, preclinical, and clinical studies, and efficiently models bulk as well as scRNA-seq data (**Fig. 1A**).

**Figure 1.**
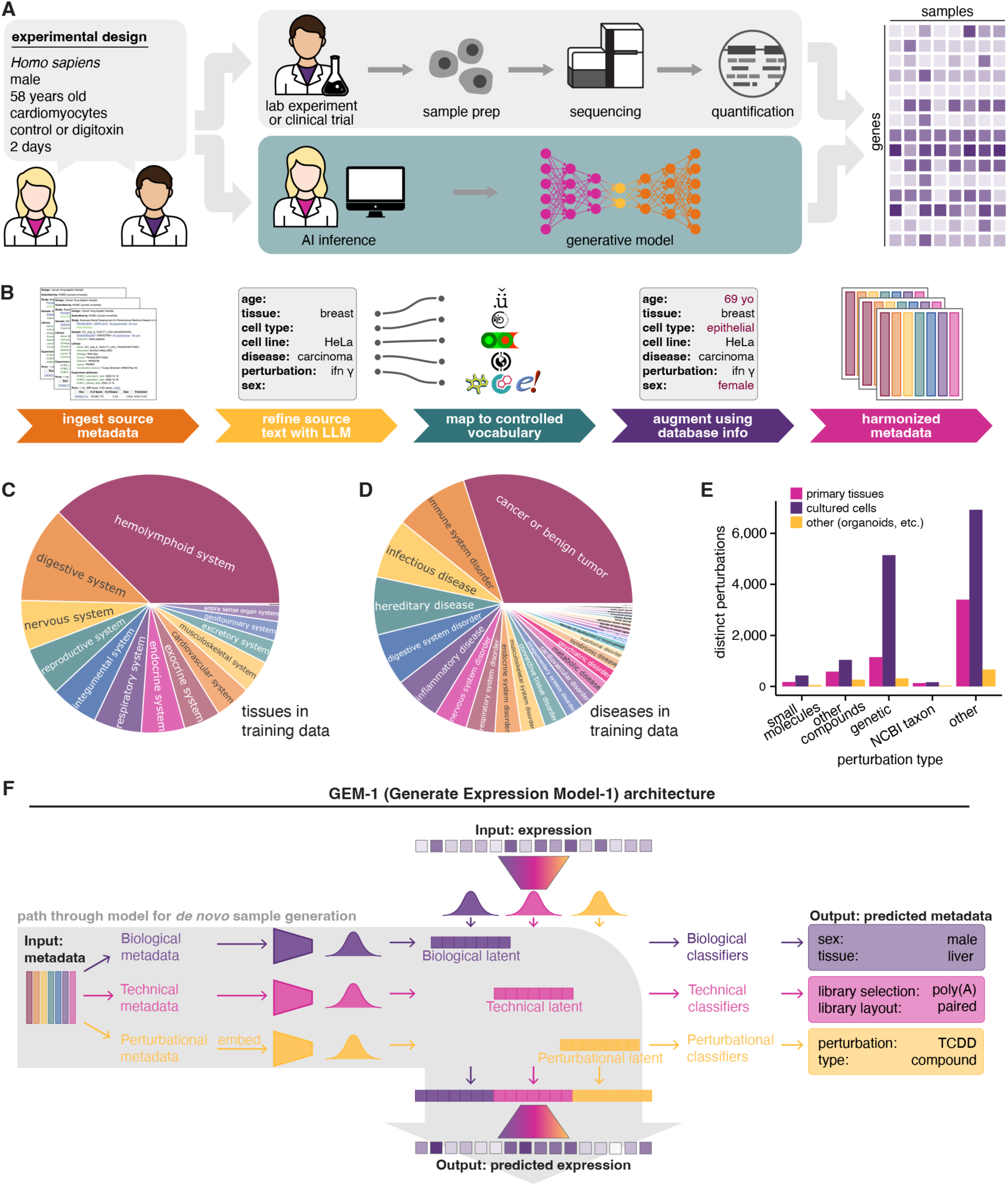
Design and construction of GEM-1. (**A**) Schematic illustrating our core problem statement: to create a generative model that mirrors the process of a scientist conducting a laboratory experiment or clinical trial and measuring resulting gene expression. (**B**) Schematic illustrating our metadata agent for processing and harmonizing source metadata. (**C**) Sunburst plot illustrating the distribution of bulk RNA-seq training data among tissue types. Plot restricted to samples from primary tissues and cell lines. (**D**) As (**C**), but for diseases. Plot restricted to samples from primary tissues (as most cell lines are cancerous). (**E**) Bar plot illustrating the numbers of distinct perturbations in bulk RNA-seq training data for different perturbation categories. “Small molecules,” compounds mapped to ChEBI identifiers; “other compounds,” compounds mapped to ChEMBL identifiers (excluding those present in ChEBI); “genetic,” perturbations to genes; “NCBI taxon,” perturbations mapped to NCBI taxa (e.g., viral infection); “other,” perturbations not included in the previous categories. (**F**) Schematic illustrating GEM-1’s architecture and path through the model for *de novo* sample generation.

## RESULTS

### Construction of a training dataset that captures the diversity of real-life experiments

We constructed a training dataset consisting of paired gene expression measurements and experimental descriptions across the NCBI SRA’s archives of RNA-seq data. These data constitute a powerful but challenging source of information for training an AI model – powerful because they are a realistic representation of the actual experiments conducted by researchers, but challenging because their very realism also renders them fragmented, incompletely described, and subject to potentially confounding technical and biological variation.

We initially focused on bulk RNA-seq data from human samples. We identified 589,542 samples from 26,680 datasets in the SRA deposited through June 30, 2024 and quantified expression of 44,592 genes in each sample, yielding a total of 470,691 samples from 24,715 datasets that passed our quality-control thresholds (details in **Methods**).

We then constructed a metadata agent to automate the process of metadata curation and harmonization in order to create descriptions of the experiments from which each RNA-seq sample was derived. This agent utilized a large language model (LLM) and biological databases to convert raw metadata from the SRA into harmonized metadata phrased with a controlled vocabulary that specified experimental parameters such as cell type, disease, and information on perturbations (**Fig. 1B**).

The resulting training dataset spanned a broad range of tissue types, with a fairly even split between data from primary tissues and cell lines. Accessible tissues such as blood were naturally most well represented in primary tissues, but the dataset had good representation of diverse other tissues as well (**Fig. 1C**, **S1A**). Cancers were the most common source of diseased primary samples, followed by immune disorders, infectious diseases, and digestive diseases (**Fig. 1D**, **S1B**). Our training dataset contained samples with over 18,000 distinct perturbations, ranging from small molecule treatment (more than 650 distinct perturbations) to pathogen infection (more than 300 distinct perturbations; **Fig. 1E**).

### GEM-1 jointly models experimental designs and gene expression results

We sought to build an AI system with an interpretable architecture that reflects the structure of biological experimentation and offers a unified framework for modeling different sources of gene expression data, including both bulk and scRNA-seq. To this end, we constructed Generate Expression Model-1 (GEM-1), a deep latent variable model (DLVM) that takes experimental metadata as input and produces gene expression experimental results as output (**Fig. 1F**). We trained GEM-1 using our dataset of gene expression measurements and paired harmonized experimental metadata.

GEM-1 partitions experimental metadata into three independent sources: biological (e.g., sex, tissue, disease), technical (e.g., library preparation method, sequencing platform), and perturbational (e.g., perturbation identity, dose, time). When possible, perturbation identity was embedded using relevant pretrained foundation models (see **Methods**). GEM-1 employs a neural network that uses this metadata and the resulting embeddings to estimate distinct latent distributions for the biological, technical, and perturbational sources. These latent distributions are then used to parameterize a final distribution over observed gene expression values. Additionally, the network includes internal classifiers that operate on samples drawn from the latent distributions to predict the most likely metadata labels (see **Methods** for a full description).

GEM-1’s interpretable architecture facilitates different model invocations to solve distinct biological problems. Our central goal–to predict the results of future gene expression experiments–is addressed by using GEM-1 for *de novo* sample generation, wherein input metadata are used to predict output expression data (**Fig. 1F**). However, GEM-1 can also be used in different modes to solve distinct problems. The separable latent generative factors used for metadata descriptions permit inference of the results of a future gene expression *conditioned on data from a previous experiment*. This mode, which we term “reference conditioning,” permits lab-informed AI experiments, such as simulating the results of a future gene knockdown in a given cell type conditioned on existing experimental data from unperturbed cells (**Fig. S1C**). GEM-1’s classifiers are similarly useful; for example, they can be used to infer sample metadata based on lab-or clinic-derived gene expression results (**Fig. S1D**).

### GEM-1 predicts future gene expression experimental results with lab-level accuracy

We constructed four distinct test datasets using holdout data (data that GEM-1 never saw during training) deposited in the SRA after our training data cutoff. We classified each dataset based upon whether its samples contained only previously observed perturbations and biological contexts, novel genetic perturbations, novel chemical perturbations, or novel biological contexts (novel indicating not observed during training; **Fig. 2A**). Interestingly, ∼95% of all holdout samples correspond to previously observed perturbations and biological contexts, indicating that our training dataset has high coverage of key features of the space of ongoing experiments.

**Figure 2.**
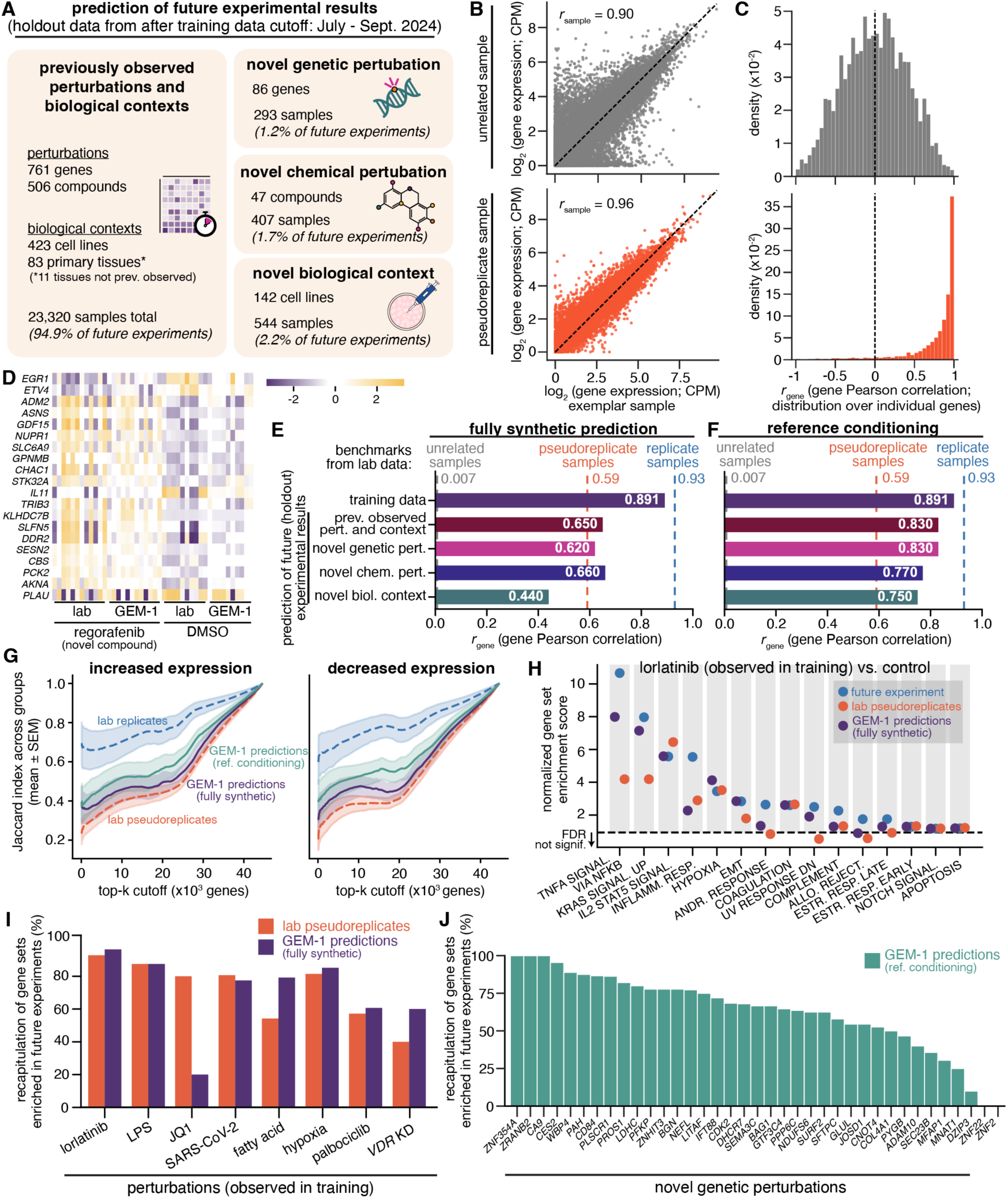
GEM-1 accurately models observed as well as novel experiments. (**A**) Distribution of holdout datasets used to test GEM-1’s ability to predict the results of future gene expression experiments. “Novel genetic perturbation,” samples containing a perturbed gene that was not perturbed in our training data; “novel chemical perturbation,” samples treated with a compound that was not present in our training data; “novel biological context,” samples arising from a cell line that was not present in our training data; “previously observed perturbations and biological contexts,” all other holdout samples (note that this includes 11 tissues that were not present in our training data; these novel tissues were not used to create a distinct holdout set given their rarity). (**B**) Top, scatter plot comparing gene expression in an exemplar sample and an unrelated, randomly chosen sample. Bottom, as top panel but for a pseudoreplicate sample (pseudoreplicate defined as a sample with matched metadata from a different study; see main text for full description). *r*sample, Pearson correlation for gene expression in the two illustrated samples. CPM, counts per million. (**C**) Top, histogram illustrating the distribution of *r*gene (gene Pearson correlation) across all genes for an exemplar sample and an unrelated, randomly chosen sample. Bottom, as top panel but for a pseudoreplicate sample. (**D**) Heat map comparing gene expression in control and regorafenib-treated glioma (LN-308) cells. Regorafenib was not observed during model training. Lab data are from a holdout lab experiment (BioProject PRJNA1138642). Plot illustrates the genes with the highest absolute log fold-changes in lab data. Data for each gene is mean centered. (**E**) Left, bar plot illustrating model performance with fully synthetic prediction across the training dataset and each of the four holdout datasets. Benchmarks from lab data are as follows: “unrelated samples,” randomly chosen samples; “pseudoreplicate samples,” samples with matched metadata from different studies; “replicate samples,” replicates with matched metadata in the same study. See **Methods** for a full description of benchmark estimation. (**F**) As (**E**), but for model performance with reference conditioning, where the model was conditioned on a control, unperturbed sample from each study and then used to generate samples with new perturbational contexts as appropriate. (**G**) Plot of Jaccard index (size of intersection divided by size of union) illustrating the overlap between the top k genes (x axis) with increased (left) or decreased (right) expression for the indicated comparisons. Analysis was performed across all studies in the holdout sets for which pseudoreplicate samples could be identified. Pseudoreplicate data were restricted to groups that had (1) available replicates to permit computation of a Jaccard index using replicates and (2) matching controls or a comparison group relative to which to define increased/decreased expression. (**H**) Dot plot illustrating gene set enrichment following lorlatinib treatment of NSCLC cells in a holdout study (BioProject PRJEB53629), a pseudoreplicate study (BioProject PRJEB50324), and AI data. Plot illustrates the most enriched Hallmark Pathways from MSigDB, identified using a hypergeometric test on log fold-change values from the holdout study. (**I**) Bar plot illustrating the percentages of gene sets that were enriched in the indicated holdout studies following the indicated perturbations and also enriched in either lab pseudoreplicate studies or fully synthetic AI predictions. Plot restricted to the eight holdout studies for which we identified matching pseudoreplicates in the training data that contained both well-defined perturbations and appropriate metadata-matched controls. (**J**) As (**I**), but for novel genetic perturbations. Plot restricted to the 37 genetic perturbations for which we identified metadata-matched control (unperturbed) and perturbed samples in holdout data.

We considered two primary statistics to assess model performance. The Pearson correlation of gene expression between two samples (*r*_sample_) is a commonly used measure of transcriptome similarity. However, because gene expression is highly structured, even unrelated samples exhibit high *r*_sample_ values (**Fig. 2B**). We therefore defined an additional statistic *r*_gene_, which tests a model’s ability to rank the expression of a given gene across samples and therefore normalizes away the baseline similarities of transcriptomes. While *r*_sample_ = 0.9 for exemplar unrelated samples, *r*_gene_ has a median of 0 (**Fig. 2C**; see **Methods** for further discussion).

We used GEM-1 to predict expression data given the metadata for each sample across all four holdout sets. Anecdotal inspection revealed concordance between transcriptome alterations in GEM-1 predictions and lab holdout data across different biological settings, such as treatment with a novel compound (**Fig. 2D**). We then quantified performance across all datasets to find that GEM-1 performance was highest on the training data, as expected, and only modestly lower on most holdout datasets (**Fig. 2E**, **S2A**). Performance was good across holdout data from the tissue types observed during training (**Fig. S1C-E**) and weakest in novel biological contexts, which is consistent with our one-hot encoding of cell lines and tissue types; in the absence of biologically informed embeddings, the model cannot accurately predict how a novel biological context alters the transcriptome.

We therefore tested how reference conditioning (**Fig. S1C**)–in this case, conditioning expression predictions for a new, perturbed sample on lab-derived expression data for a control, unperturbed sample in the study–affected GEM-1 performance. Adding reference conditioning yielded marked performance improvements, with accuracy on all holdout sets approaching that observed for training data, including for novel biological contexts (**Fig. 2F**, **S2B**).

We then compared GEM-1 accuracy to best-possible performance benchmarks established from lab data. We considered two benchmarks, replicates and pseudoreplicates. Replicates are defined in the traditional sense, as samples in the same study with matched metadata, while pseudoreplicates are samples in different studies with matched metadata (e.g., if the same gene is knocked down in the same cell line, but in different labs). We term these pseudoreplicates because our definition ensures that two pseudoreplicates are quite similar but nonetheless likely not identical; for example, unrecorded culture conditions could differ.

We estimated replicate- and pseudoreplicate-level reproducibility across our training and holdout datasets to find that GEM-1 exhibited pseudoreplicate-level performance when used in fully synthetic mode (“fully synthetic” meaning using only the pretrained model, without the additional information provided by reference conditioning) in all settings except for novel biological contexts, rising to performance that exceeded pseudoreplicate reproducibility across all holdout sets when used with reference conditioning (**Fig. 2E-F**, **S2A-B**).

GEM-1 performed well at future experiment prediction relative to lab benchmarks across multiple axes. We identified holdout studies with matched pseudoreplicates within the training data and assessed the extent to which GEM-1 accurately predicted holdout experimental results relative to the overlap between training data pseudoreplicates and holdout experiments. GEM-1 predictions effectively recapitulated hierarchies of differential gene expression, with performance exceeding lab pseudoreplicates (**Fig. 2G**), and GEM-1-predicted differential gene expression was enriched for the specific biological pathways observed in holdout perturbational experiments (**Fig. 2H-I**). We extended these analyses to novel genetic and chemical perturbations to find that GEM-1 correctly predicted over half of gene sets enriched in future experiments in fully synthetic mode, rising to 63% for genetic perturbation and 70% for chemical perturbations when reference conditioning was used (**Fig. 2J**, **S3**).

Overall, these data demonstrate that GEM-1 can successfully predict molecular phenotypes of gene expression experiments from experimental descriptions, including in the setting of previously unseen perturbations.

### An interpretable model architecture enables biological feature inference

We next assessed the interpretability of core features of GEM-1. We visualized GEM-1’s embeddings by passing gene expression counts from the training data through GEM-1’s expression encoder and performing dimensionality reduction with the UMAP algorithm^24^. Visual inspection of the resulting plots confirmed that samples clustered by biological parameters in the biological, but not technical, embedding and by technical parameters in the technical, but not biological, embedding (**Fig. 3A**).

**Figure 3.**
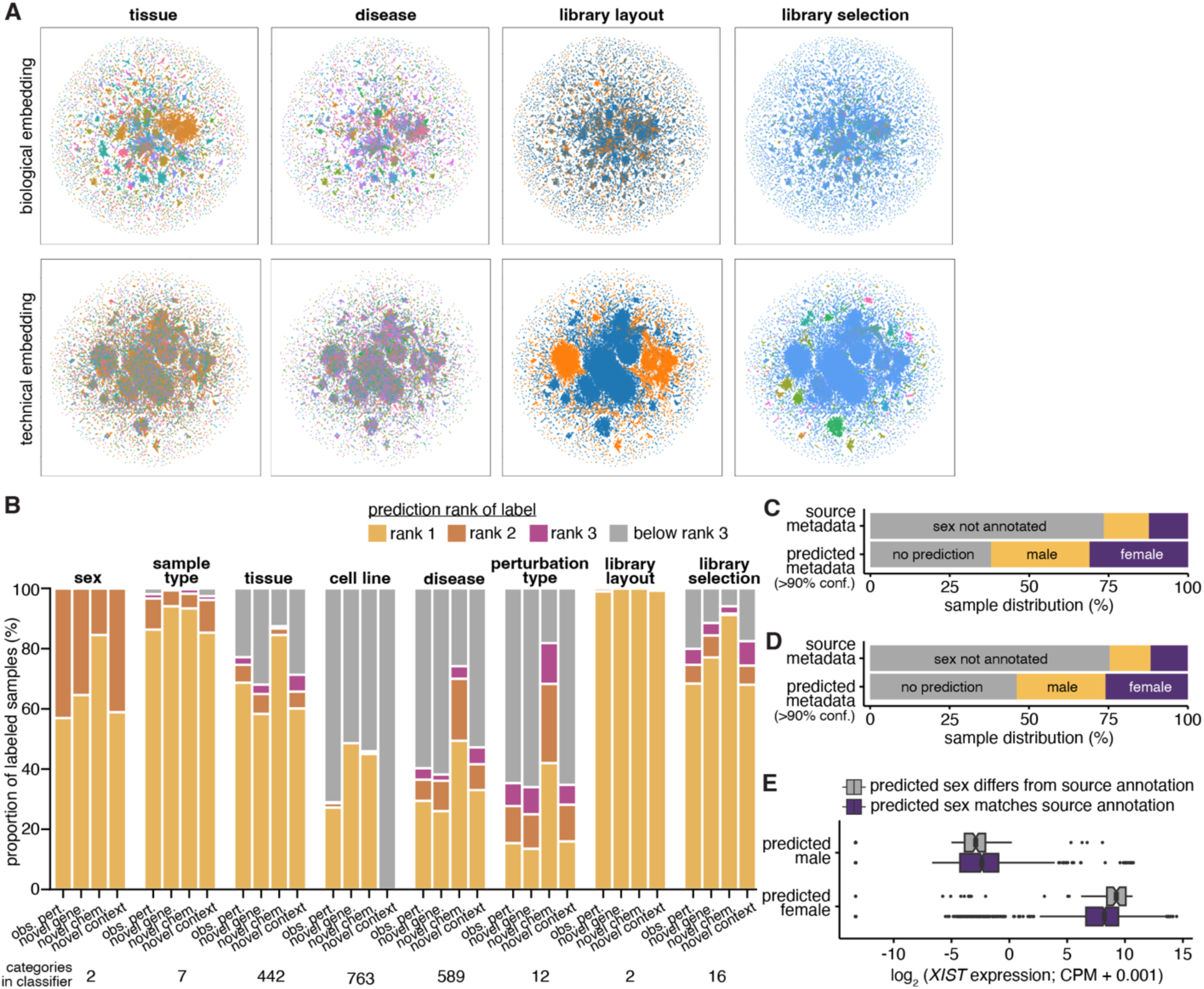
Learned embeddings and classifiers distinguish key biological features. (**A**) Top row, UMAPs illustrating projection of the training data using GEM-1’s learned biological embedding. Points are samples, colored by tissue, disease, library layout, or library selection method. Bottom row, as top row, but using GEM-1’s learned technical embedding. (**B**) Bar plot illustrating the ranked prediction performance of GEM-1’s classifiers on each holdout set. The bars show the percentages of samples for which the correct metadata label was predicted at a given rank. Each classifier analysis was restricted to samples annotated with the relevant metadata. (**C**) Distribution of samples in the training data with sex annotations available and/or predicted with >90% confidence. (**D**) As (**C**), but for samples in the holdout data. (**E**) Box plot of *XIST* expression in holdout samples categorized as indicated. Boxes show the median and lower and upper quartiles of data. Whiskers extend from the interquartile range (IQR) to the largest value no further than 1.5 * IQR. Points beyond the whiskers are plotted individually. Notches indicate 90% confidence interval for the median. CPM, counts per million.

GEM-1’s classifiers are useful for both model training and inference. In addition to encouraging linear separability of the latent spaces, the classifiers permit inference of metadata conditioned on input, lab-derived expression data (**Fig. S1D**). We estimated classifier accuracy by predicting biological and technical features of each holdout dataset given the expression data alone and compared the resulting predictions to metadata annotations. Most classifiers performed with high accuracy; for example, the top-ranked tissue prediction (out of 442 tissue categories) corresponded to the metadata label for ∼60-80% of samples across the holdout datasets (**Fig. 3B**).

We performed a detailed investigation of differences in predicted and annotated metadata for sex. Sex was annotated in ∼25% of samples in the training as well as holdout datasets, while GEM-1 predicted sex with >90% confidence in the majority of samples in both datasets (**Fig. 3C-D**). We estimated the quality of sex inference by examining expression of *XIST*, the gene responsible for X chromosome inactivation in female cells. *XIST* expression was strongly bimodal between predicted male and female samples irrespective of whether the prediction matched or differed from the annotation (**Fig. 3E**), consistent with prediction accuracy exceeding source annotation accuracy. These analyses indicate that GEM-1’s classifiers can be used for filling in missing metadata as well as detecting metadata inaccuracies.

### GEM-1 is readily extensible to single-cell data

We next tested GEM-1’s utility for modeling scRNA-seq data. We trained GEM-1 using publicly available, harmonized data from 41,474,950 unique cells, representing 669 distinct cell types and 259 distinct tissues^16,25^. We took advantage of the two distinct data releases^26,27^ from the Tabula Sapiens consortium to construct a holdout dataset for testing model performance. We included the first data release in our training data and used the second data release, which included 4 additional organs and 9 additional study participants, as a holdout dataset.

We visualized GEM-1’s embeddings of the holdout dataset by passing expression data through the expression encoder and performed dimensionality reduction with the UMAP algorithm^24^. We compared to embeddings from scGPT^19^, a widely used, pretrained model of scRNA-seq, for a reference point. GEM-1’s biological embedding yielded a stronger separation of cells by cell type than did GEM-1’s technical embedding, as expected, and was qualitatively similar to scGPT’s embedding (**Fig. 4A**, **S4A**).

**Figure 4.**
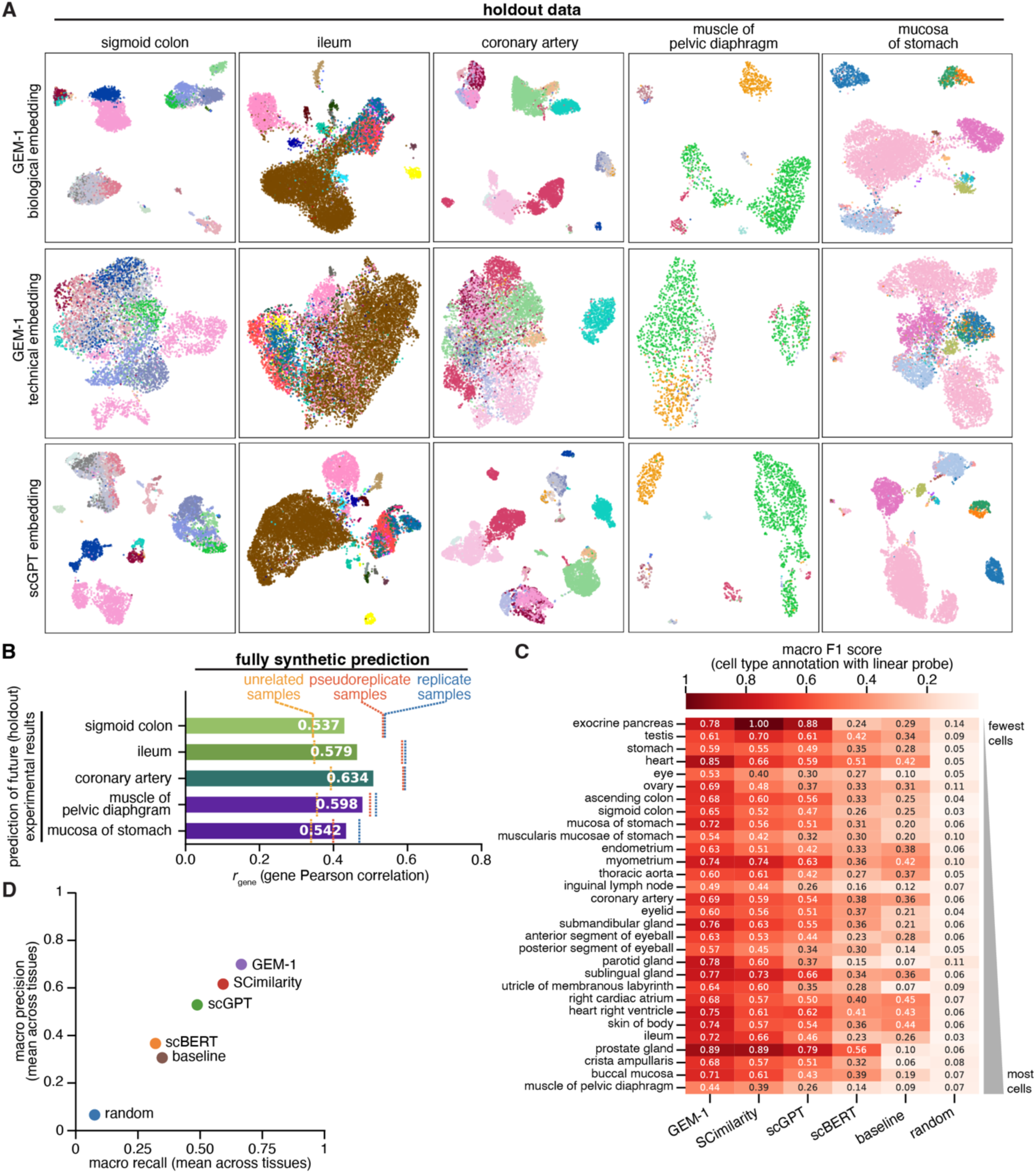
GEM-1 is readily extensible to model single-cell data. (**A**) Top row, UMAPs illustrating projection of holdout data from the indicated tissues using GEM-1’s learned biological embedding. Middle row, as top row, but using GEM-1’s learned technical embedding. Bottom row, as top row, but using embeddings from scGPT. Points are cells, colored by cell type; see **Fig. S4A** for legends showing the mapping between colors and cell types. (**B**) Bar plot illustrating model performance for holdout data from the indicated tissues (fully synthetic prediction). Benchmarks from lab data are as follows: “unrelated samples,” randomly chosen samples; “pseudoreplicate samples,” samples from the same tissue and cell type, but from different donors; “replicate samples,” samples from the same tissue, cell type, and donors. These benchmarks are computed on a per-tissue level due to the structure of the Tabula Sapiens dataset (c.f., the benchmarks for bulk RNA-seq). See **Methods** for a full description of benchmark estimation. (**C**) Heat map illustrating the performance of different models for cell type annotation on holdout data (not seen by any evaluated model during training). For each model, cell-level embeddings were extracted from the pretrained weights and used to train a linear probe classifier. The probe was implemented as a logistic regression model trained with an 80/20 train/test split within each tissue, stratified by cell type where possible to maintain class balance. “baseline” indicates GEM-1’s performance when predictions were made using the pretrained cell type classifier alone, without a linear probe. “random” indicates a baseline in which each cell was assigned a randomly generated embedding of the same dimensionality as the learned representations. The macro F1 score for a given tissue was computed by taking an unweighted average of the F1 score across all tissue-donor combinations. The F1 score itself was calculated as the harmonic mean of precision (the proportion of predicted cells that were correctly assigned to a type) and recall (the proportion of true cells of a type that were successfully identified). Cell number corresponds to the number of cells in the test split (holdout data) for each tissue. (**D**) Scatter plot illustrating the precision and recall, averaged across all tissues in the holdout data, for the indicated models when used for cell type annotation with a linear probe.

We assessed GEM-1’s performance on two tasks, our core goal of data generation and the common scRNA-seq problem of cell type annotation. We used GEM-1 to generate transcriptomes using metadata from the holdout dataset as input and estimated lab reproducibility benchmarks using replicates and pseudoreplicates, defined as matched cells from identical or different donors, respectively. As these pseudoreplicates were generated by the same consortium, this definition is an overly optimistic estimate of experimental reproducibility across labs, versus the more realistic estimate for bulk RNA-seq. The similarities between GEM-1-predicted transcriptomes and holdout data were fairly constant across tissues, while lab reproducibility exhibited greater variation (**Fig. 4B**).

GEM-1’s internal classifiers offer a unifying approach to metadata annotation for both bulk and single-cell data. We compared GEM-1’s ability to annotate cell types from expression data to cell type annotation with several widely used, pretrained deep learning models^17,19,28^.

Following the standard linear probe framework^29^, we used holdout expression data as input to each pretrained model, extracted cell-level embeddings, trained a linear classifier on these frozen embeddings, and applied the probe to predict cell type labels. GEM-1, scGPT, and SCimilarity each exhibited relatively comparable precision and recall, with GEM-1’s performance being modestly better on most tissues (**Fig. 4C-D**, **S4B**).

### AI-generated cohorts capture key biological phenomena observed in large, holdout clinical studies

We evaluated GEM-1’s utility for generating data from large clinical cohorts, which are essential for translational studies but difficult to assemble and subject to strong regulatory constraints. We studied three biological contexts: healthy tissues, autoimmune disease, and oncology.

We used GEM-1 to generate expression data from 30 distinct healthy tissues for a total of 5,300 samples. This SYNTH-TEx dataset is comparable to transcriptome data from the Genotype-Tissue Expression (GTEx) project^30^, which we excluded from our training data. We overlaid and visualized data from SYNTH-TEx and GTEx with dimensionality reduction to find that samples clustered by tissue (**Fig. 5A**), with GEM-1 and the holdout clinical data closely overlaid in most cases (**Fig. 5B**).

**Figure 5.**
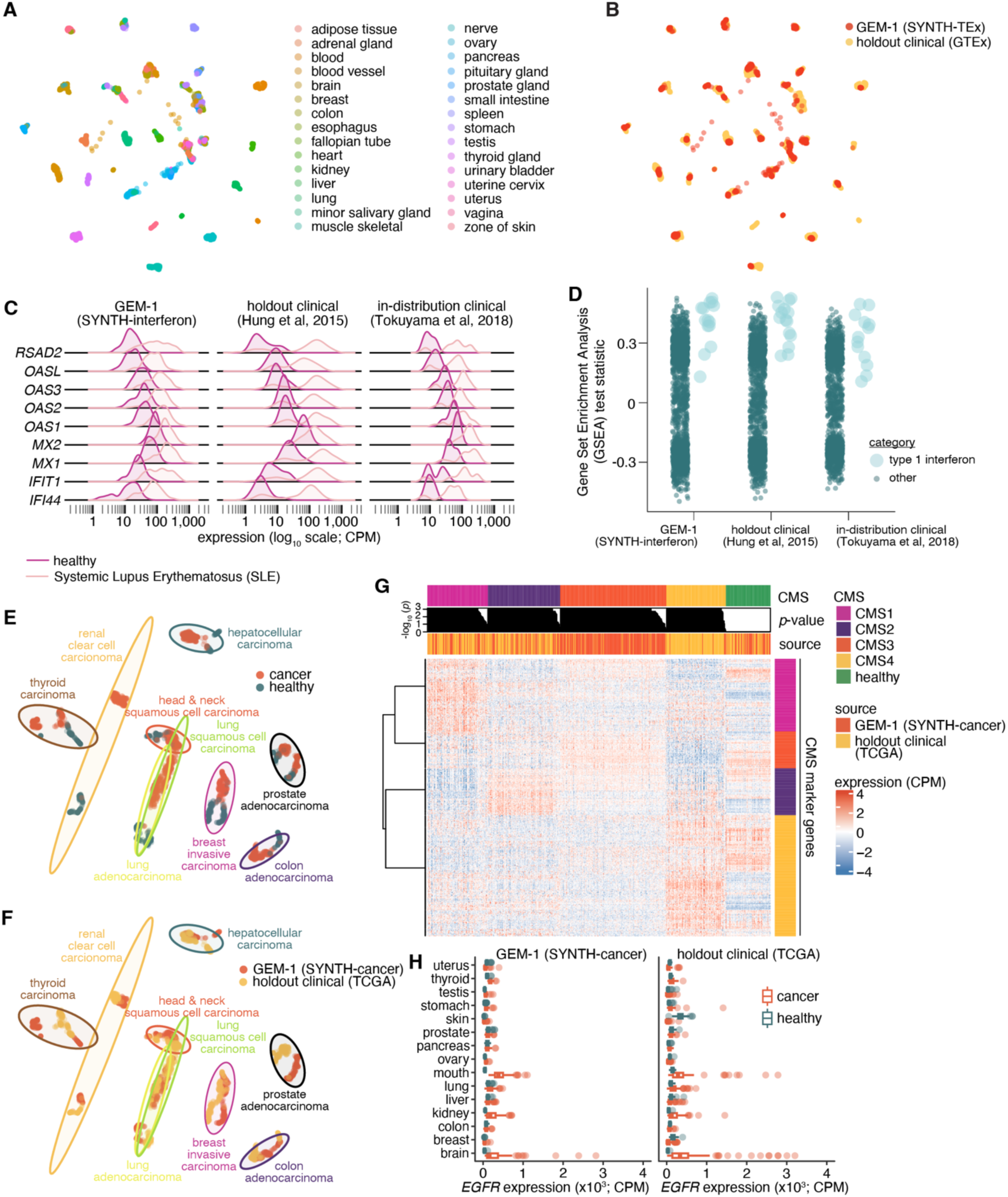
AI-generated cohorts capture key biological features of clinical cohorts. (**A**) UMAP illustrating AI (SYNTH-TEx) and holdout clinical (GTEx) data together. Points are samples, colored by tissue. UMAP created by calculating embeddings on GTEx samples that were used to project SYNTH-TEx into the same latent space (**B**) As (**A**), but colored by sample origin (SYNTH-TEx or GTEx). (**C**) Expression of the indicated IFN-responsive genes in healthy and SLE samples from AI (SYNTH-interferon) and holdout as well as previously observed (in-distribution) clinical studies. (**D**) Dot plot showing the distributions of GSEA test statistics from all gene sets in Connectivity Map (CMAP) cytokine perturbations (i.e. “genetic reagents”) that are or are not associated with a Type 1 interferon response in AI and clinical studies. (**E**) UMAP illustrating AI (SYNTH-cancer) and holdout clinical (TCGA) data together. Points are samples, colored by disease status. UMAP computed using AI and clinical data together. Plot restricted to 100 randomly chosen samples from each data source (AI or clinical)-disease status combination for simplicity of presentation. (**F**) As (**E**), but colored by data source (SYNTH-cancer or TCGA). (**G**) Heat map illustrating expression of CMS marker genes across AI and clinical data. Samples and genes were ordered by CMS group and then clustered within each group (unsupervised). (**H**) *EGFR* expression in healthy and cancer samples across major organs and cancer indications. Sample set sizes for both data generation types match those of the TCGA study. Boxes show the median and lower and upper quartiles of data. Whiskers extend from the interquartile range (IQR) to the largest value no further than 1.5 * IQR. Points beyond the whiskers are plotted individually.

We next generated expression data corresponding to blood samples taken from a cohort of individuals who were healthy (n = 100) or diagnosed with an autoimmune disease, Systemic Lupus Erythematosus (SLE; n = 100). We compared this SYNTH-interferon dataset to data from two clinical SLE cohorts, one holdout^31^ and one observed during training^32^. Dysregulation of Type 1 interferon is a key feature and driver of SLE pathology^33^. Accordingly, select Type 1 interferon-stimulated genes (ISGs) were upregulated in SLE relative to healthy samples across SYNTH-interferon as well as the two clinical cohorts (**Fig. 5C**), and a transcriptome-wide analysis of Type 1 interferon-related gene sets showed specific enrichment in SLE versus healthy samples as well (**Fig. 5D**).

Finally, we generated expression data from 27 cancer types, as well as normal control samples when relevant, for a total of 10,523 samples. We designed this SYNTH-cancer dataset to mimic the structure of The Cancer Genome Atlas (TCGA), which we excluded from our training data. SYNTH-cancer and holdout TCGA data exhibited similar structures, with samples clustering by tissue of origin and then healthy versus cancer (**Fig. 5E-F**). SYNTH-cancer samples exhibited expected molecular signatures at both whole-transcriptome and single-gene levels. For example, SYNTH-cancer samples conformed to commonly used molecular subtyping schemes, such as the Consensus Molecular Subtypes (CMS) for colorectal carcinoma (**Fig. 5G**), and the expression distribution of key therapeutic targets such as *EGFR* was similar across cancers between SYNTH-cancer and holdout clinical data (**Fig. 5H**).

Each of these three datasets (SYNTH-Tex, SYNTH-interferon, and SYNTH-cancer), along with additional analyses and comparisons to holdout clinical studies, is publicly available (https://app.synthesize.bio/datasets).

## DISCUSSION

Here, we provide evidence of feasibility of our central goal – to overcome fundamental limitations in biological experimentation and clinical trials by generating essential data with an AI system. Our approach explicitly reflects the nature of biological experimentation, wherein our deep latent variable model (DLVM) deconvolves gene expression data into its core biological, perturbational, and technical components to provide an interpretable framework for flexible *in silico* experimentation.

A key innovation is GEM-1’s ability to support nuanced conditional sampling, which we term “reference conditioning.” This allows researchers to, for example, simulate a novel drug’s effect on a specific patient’s tissue sample by combining latent factors inferred from observed data with factors sampled from metadata-conditioned priors. This “mix-and-match” capability is a direct result of the evidence lower bound (ELBO) objective, which balances two complementary goals. The reconstruction term encourages a low-entropy variational posterior (*Q*) to capture the unique fingerprint of a specific sample for high-fidelity reconstruction, while the KL divergence terms ensure that this specific representation lies within a broader, high-entropy prior (*Pr*) that models the full heterogeneity of the population. This interplay between a specific posterior and a general prior is what grants researchers the unique ability to generate data that is either precisely faithful to a reference sample or realistically diverse for population-level simulations.

Our study also illustrates several important concepts for biology-focused AI models. First, much as LLMs made rapid progress by modeling virtually all available text rather than focusing on more narrowly defined corpuses, we found explicitly modeling the wide diversity of experiments represented by the NCBI SRA (rather than focusing on a narrower dataset) to be similarly fruitful. Second, our study highlights the importance of metadata curation and training data quality for AI models. Our rich metadata resource also provided unexpected insights. For example, ∼95% of future experiments in our holdout data (on and after July 1, 2024) represented perturbations and/or biological contexts that we had previously observed in training data (from before July 1, 2024; **Fig. 2A**). Third, an interpretable model architecture facilitates subsequent model utilization in real-world settings, whether checking sample provenance (**Fig. 3B**) or simulating large patient cohorts (**Fig. 5**).

Despite the advances described in this manuscript, there are several limitations to note. First, we focused on human data alone, although our model is readily extensible to a multi-species context. Second, our metadata schema captures many, but not all, features of an experiment; we expect GEM-1’s performance to further improve as the input metadata is more fully specified. Third, GEM-1’s accuracy in different settings reflects its training data distribution. It therefore provides a more realistic representation of day-to-day experiments conducted by typical researchers for bulk RNA-seq than for scRNA-seq, simply because most scRNA-seq data has been generated by consortia primarily focused on large-scale data generation (a powerful but atypical setting). Fourth, while our models encompass a wide array of gene expression experiments, we did not include extensive Perturb-seq data during training. Despite being a valuable resource for AI model development^20,34,35^, Perturb-seq data presents unique challenges, including technical artifacts and non-specific responses^20,36–39^, which require focused resolution for this data type that goes beyond the scope of our initial study.

Finally, it is important to note that we expect that our model will not stand in isolation but rather both benefit from and contribute to the growing ecosystem of deep learning-powered tools in biomedicine. GEM-1 performed well even when confronted with novel genetic as well as chemical perturbations (**Fig. 2** and **S2**), which we conjecture reflects not just the utility of our model but also the strengths of the biologically informed embeddings that we relied upon to represent these perturbations. We speculate that our model’s ability to effectively generalize outside of its training data will continue to grow in concert with improvements in embeddings.

## METHODS

### Training dataset construction

**Bulk RNA-seq processing.** Bulk RNA-seq data was obtained from the NCBI Sequence Read Archive (SRA)^12^. Samples were selected by querying for all human transcriptomic samples sequenced on an Illumina platform without size fractionation as the library selection method.

The FASTQ files from all samples meeting these criteria were pulled using a custom Nextflow pipeline that utilized the SRA toolkit to fetch the files and subsequently quantified counts using Kallisto^40^ with the Ensembl 111 gene annotation^41^. Kallisto was called with default parameters (estimate fragment length of 200 and sd of 20 for single-end data; not applicable for paired-end data). For data used in training, we restricted to protein-coding genes, long non-coding RNAs, and immunoglobulin-related genes, as well as to samples that passed basic quality-control checks (less than 80% read duplication rate, at least 50% of reads mapped, and at least 10,000 genes with non-zero counts). All data deposited in the SRA through June 30, 2024 was ingested, processed, and used for model training assuming that it passed the described quality-control checks.

**Single-cell RNA-seq processing.** Single-cell RNA-seq data was obtained from the CZI CellxGene^16^ and the scPerturb^25^ databases. We used the entire July 2024 version of CellxGene data, aligned with the Ensembl 110 gene annotation^41^. We also used 10 datasets from scPerturb, which were provided as reformatted count matrices pulled from GEO: JoungZhang2023_atlas, ReplogleWeissman2022_K562_gwps, ReplogleWeissman2022_rpe1, ReplogleWeissman2022_K562_essential, SrivatsanTrapnell2020_sciplex3, TianKampmann2019_iPSC, ZhaoSims2021, AdamsonWeissman2016_GSM2406681_10X010, AdamsonWeissman2016_GSM2406675_10X001, AdamsonWeissman2016_GSM2406677_10X005. For data used in training, we restricted to protein-coding genes, long non-coding RNAs, and immunoglobulin-related genes, as well as to cells that had at least 200 genes with non-zero counts.

**Metadata agent.** Metadata was pulled as available from all sources and harmonized using an agent that accomplished the following tasks. The metadata was standardized as much as possible into chosen feature columns (age, sex, tissue, cell type, disease, etc.) from source information using a Large Language Model (LLM; Claude 3.5 Sonnet). The LLM was given free text metadata at both the study and sample levels and prompted to label individual samples.

This output was then programmatically cleaned using observation-based rules. Perturbations, tissues, cell types, cell lines, and diseases were then mapped to corresponding ontologies or controlled vocabularies. The text2term^42^ package was utilized to assist with mapping cleaned text to ontologies; synonyms were additionally used to map cleaned text to controlled vocabularies. The cell type labels were mapped to the cell ontology^43^, cell line labels were mapped to Cellosaurus^44^, tissue labels were mapped to the Uberon ontology^45^, and disease labels were mapped to the Mondo ontology^46^. Within the perturbation labels, genes were mapped to Ensembl stable gene identifiers^41^, small compounds were mapped to the Chemical Entities of Biological Interest (ChEBI) ontology^47^, other bioactive molecules were mapped to ChEMBL^48^ (e.g., compounds that did not map to ChEBI identifiers), and viruses and bacteria were mapped to the NCBI Taxonomy ontology^49^. Perturbations that did not fall under one of these categories were retained as standardized text labels.

### Test dataset construction

**Construction of holdout datasets.** For bulk RNA-seq, holdout datasets that were used to assess our model’s ability to predict future experimental results were constructed using data deposited in the SRA during the date range July 1, 2024 through September 30, 2024. Three distinct subsets were then created from this “out-of-time” holdout dataset by identifying SRA studies that contained a novel genetic perturbation, a novel chemical perturbation, or a novel biological context, where “novel” means not contained in the training dataset. The novel biological context subset was restricted to datasets containing novel cell lines; because there were relatively few studies that contained novel primary tissues (n = 11 tissues in the out-of-time dataset that were not previously observed during training), we did not create a subset specific to novel primary tissues. All holdout samples that did not fall into one of those three subsets were assigned to a fourth subset. See **Fig. 2A** for an illustration of these four subsets and associated information on sample number, etc.

For single-cell RNA-seq, a holdout dataset was constructed based on the Tabula Sapiens dataset, which was generated by the Tabula Sapiens consortium^26,27^. Tabula Sapiens data was released by the consortium in two versions. The first version^26^ (v1) contained data from 24 organs from 15 human study participants. The second version^27^ (v2) was expanded to include 4 additional organs and 9 additional study participants. The first version of Tabula Sapiens was included in our training data, while data from the 9 new donors included in version two of Tabula Sapiens were used as a holdout set for testing model performance.

**Identification of replicate and pseudoreplicate lab studies and establishment of performance benchmarks** For bulk RNA-seq analyses, within-study replicates were defined as pairs of samples originating from the same study that shared identical experimental and biological metadata (e.g., perturbation type, cell line, tissue, etc.). For this benchmark, 200 distinct replicate pairs across 61 studies were randomly selected from the holdout data (see below for additional details). Pseudoreplicates were identified by finding metadata-matched samples across studies. We defined pseudoreplicates as samples in different studies which had exactly matched metadata across 12 fields describing the experiment, including tissue, disease, perturbation ontology identifier, cell line, and demographic data (e.g., age, sex, and race). We performed this metadata matching in two contexts: (1) within the training data, in which we identified 10,000 matched sample pairs across 2,511 studies, and (2) between the training and holdout data, yielding 1,662 matched sample pairs in 172 study pairs. In both cases, we required that a given study only be used in one study-pair match. These sets of replicates and pseudoreplicates from lab data were used to estimate corresponding lab performance benchmarks illustrated in **Fig. 2**. For the illustrations of gene and sample Pearson correlations (**Fig. 2B-C**), we utilized an exemplar sample from BioProject PRJNA1161150, an unrelated sample from BioProject PRJEB53629, and a pseudoreplicate from BioProject PRJDB4300.

For single-cell RNA-seq, we defined replicates and pseudoreplicates using the Tabula Sapiens data. Replicates were defined as cells with matching metadata for tissue, cell type, cell line and donor ID. Pseudoreplicates were defined as cells with matching metadata for tissue, cell type and cell line, but not matching donor ID. These sets of replicates and pseudoreplicates from lab data were used to estimate corresponding lab performance benchmarks illustrated in **Fig. 4**.

### Model evaluation

**Evaluation metrics.** For bulk RNA-seq, model performance was evaluated using two Pearson correlation metrics (*r*_sample_ and *r*_gene_) computed using log-normalized expression data (in units of CPM). These two Pearson correlation metrics (*r*_sample_ and *r*_gene_) are defined in detail below in terms of comparing a lab study to an AI-generated study; however, they can be identically computed for the case of comparing data between two lab studies.

The sample Pearson correlation (*r*_sample_) assesses the fidelity with which an AI-predicted gene expression profile recapitulates the overall expression profile of a given lab sample. It is initially computed for a given sample across all genes by calculating the Pearson correlation coefficient between the vector of predicted and observed expression values across all genes for a given sample. The mean across all per-sample correlation coefficients is then taken to yield a single statistic for all samples under consideration.

The gene Pearson correlation (*r*_gene_), in contrast, assesses a model’s prediction accuracy for a given gene across all samples. In essence, it tests the model’s ability to rank the expression of a given gene across samples. It is initially computed for a given gene across all samples by calculating the Pearson correlation coefficient between the gene’s vector of predicted and observed expression values across all samples under consideration. This per-gene metric is then used to compute a variance-weighted average of per-gene correlation coefficients across all genes to yield a single statistic across all samples under consideration.

This variance weighting (with the weighted variance for a given gene defined as that gene’s correlation multiplied by its observed variance across all samples under consideration and then divided by the sum of per-gene variances across all genes) gives greater weight to genes with higher variance in the observed data, helping to ensure that the metric is focused on true biological signal rather than technical noise.

While both metrics are informative, the gene Pearson correlation (*r*_gene_) provides a more stringent evaluation of performance. The sample Pearson correlation typically remains high even between unrelated samples because gene expression profiles have a highly structured nature which is shared between biologically unrelated samples. As illustrated in **Fig. 2B-C**, this phenomenon is obvious upon comparing metadata-matched samples to unrelated samples. In the case of unrelated samples, the sample Pearson correlation is high, whereas the gene Pearson correlation approaches zero. The gene Pearson correlation is therefore a more accurate representation of the relationship between samples that is largely not influenced by the strong structures and baseline similarities between gene expression profiles.

For single-cell RNA-seq, evaluation metrics used to assess model performance on the holdout data (**Fig. 4B**) were computed on a per-sample basis and then averaged to obtain scores for each tissue type. These metrics (*r*_sample_ and mean squared error) are defined in detail below in terms of comparing a lab study to an AI-generated study; however, they can be identically computed for the case of comparing data between two lab studies.

We computed evaluation metrics using two vectors of log-normalized expression data (in units of CPM, with a pseudocount of 1): a lab-observed expression profile, denoted *lab_vector*, and a model-predicted profile, denoted *pred_vector*. *lab_vector* contained the log-normalized expression values with a pseudocount of 1 (“log1p”) for all genes, providing a standardized representation of observed expression. The *pred_vector* contained the corresponding log1p-normalized predictions, which were generated using GEM-1 conditioned on the sample’s metadata. The metadata used to generate *pred_vector* encompassed a broad set of features, including biological variables (sex, cell line ontology identifier, cell type ontology identifier, disease ontology identifier, tissue ontology identifier, sample type, age in years, ethnicity, race, genotype, and developmental stage), technical variables (library selection, library layout, platform, study, modality, and subject identifier), and perturbation variables (perturbation ontology identifier, perturbation time, perturbation dose, and perturbation type).

Two complementary metrics were used to assess predictive performance. The first was the sample Pearson correlation coefficient (*r*_sample_), which quantifies the similarity between *lab_vector* and *pred_vector* across all genes in a given sample. This coefficient directly reflects the fidelity with which the predicted expression profile recapitulates the overall observed profile. After computing the Pearson correlation for each sample, the values were averaged across all samples within a tissue type to yield a tissue-level summary statistic. The second metric was the mean squared error (MSE), which captured the magnitude of deviation between predicted and observed gene expression. For each sample, MSE was defined as the average of the squared differences between *lab_vector* and *pred_vector* across all genes, providing a measure of absolute prediction error. As with the correlation metric, MSE values were first computed per sample and then averaged across all samples within each tissue type to obtain tissue-level estimates. The gene Pearson correlation (*r*_sample_) that served as an important metric for model performance for bulk RNA-seq is less suitable for scRNA-seq data because the sparsity and prevalence of dropout events that typify scRNA-seq data cause many gene expression values to be zero, inflating correlations and obscuring meaningful biological signals.

**Performance benchmarks for lab experiments.** The performance benchmarks for different lab samples (**Fig. 2E-F**) were computed as follows. The unrelated samples benchmark was calculated by randomly shuffling 100 holdout studies into two halves and computing *r*_sample_ and *r*_gene_ between these two unrelated sets. The replicates benchmark was computed by pairing true replicate samples from within the same study and experimental condition; 200 distinct replicate pairs across 61 studies were randomly selected from those studies where there were enough samples to create replicate groups with at least 2 samples. The pseudoreplicates benchmark was computed using pseudoreplicate pairs defined as specified above, calculated using the union of the within-training data pseudoreplicates and between holdout data and training data pseudoreplicates (i.e., using all available pseudoreplicates).

For benchmarking single-cell RNA-seq performance (**Fig. 4B**), vertical dashed lines in the figures represent baseline scores derived from three control settings: pseudoreplicates, replicates, and unrelated samples. In the pseudoreplicate setting, predicted expression profiles (*pred_vector*) were obtained by averaging gene expression values across all samples within a shared metadata grouping defined by cell line ontology identifier, cell type ontology identifier, and disease ontology identifier, without regard to subject identity. Replicates were constructed in the same way, except restricted to samples derived from the same subject identifier, thereby capturing within-subject reproducibility. In contrast, the unrelated sample baseline was defined by taking *pred_vector* as the expression profile of a randomly selected training cell, providing a lower-bound benchmark for model fidelity. In each case, the observed expression profile (*lab_vector*) was defined as the log1p-normalized expression of the full gene vocabulary for the holdout sample. Performance was assessed using the same per-sample metrics described above: the mean squared error (MSE), defined as the average squared difference between *lab_vector* and *pred_vector*, and *r*_sample_, computed as the correlation between *lab_vector* and *pred_vector* across all genes. Baseline values were then averaged across samples to yield the dashed line benchmarks shown in the evaluation plots.

**Differential gene expression analysis.** The Jaccard analysis comparing genes with increased or decreased expression (**Fig. 2G**) was performed as follows. For each holdout sample for which we identified a pseudoreplicate in the training data, we generated a corresponding AI sample using the relevant input metadata. We then computed log fold-changes in gene expression (calculated against a control group matched on technical and biological metadata, falling back to a relaxed match requiring a core set of biological keys – sex, tissue, and cell line – when a direct control was unavailable) to rank the genes. When generating AI samples for fully synthetic predictions, we did not require matched controls to be present; when generating AI samples for reference-conditioned predictions, we did require the presence of matches between samples to controls on technical metadata such as study ID as well as biological metadata and perturbation time. The Jaccard index was then calculated using the resulting vectors of log fold-changes at different k intervals across four pairings: (i) a holdout sample comparison and its corresponding fully-synthetic AI prediction, (ii) a holdout sample comparison and its corresponding reference-conditioned AI prediction, (iii) a holdout sample comparison and its pseudoreplicate comparison, and (iv) biological replicates from the same experimental condition from holdout data.

**Gene set enrichment analysis.** We identified statistically enriched biological pathways in lab-and AI-derived data (**Fig. 2H-J**, **Fig. S3**) using gene set enrichment analysis, performed using the Hallmark Gene Set from the Molecular Signatures Database (MSigDB)^50^. For each experimental condition (e.g., a specific perturbation within a given cell line), genes were first ranked by their absolute log-fold change relative to control samples, and the top 100 most-altered genes were selected. Then, a one-sided hypergeometric test was used to determine if any of the hallmark gene sets were over-represented in this list of 100 genes. This test calculates the probability of observing the given overlap between the subset of genes and a hallmark pathway by random chance, using a background set of all annotated genes.

The resulting *p*-value for each pathway was transformed into a standardized Normalized Enrichment Score (NES) by taking the negative base-10 logarithm of the *p*-value to quantify the strength of enrichment. Finally, to correct for multiple hypothesis testing across all pathways, *p*-values were adjusted using the Benjamini-Hochberg procedure to control the False Discovery Rate (FDR). Hallmark pathways with FDR < 0.25 were considered significantly enriched.

**Embedding analysis.** For bulk RNA-seq, we visualized the data by computing technical and biological embeddings (**Fig. 3A**) from a random subset of 400,000 samples from the training set. Embeddings were first generated by passing observed gene expression values (counts) through our expression encoder. We then reduced their dimensionality for visualization using the UMAP algorithm^24^, implemented with the umap-learn library (https://github.com/lmcinnes/umap), and visualized the results with the matplotlib and seaborn packages in Python.

For scRNA-seq, we visualized the data by computing technical and biological embeddings of GEM-1 (**Fig. 4A**, top and middle rows) for samples from the holdout dataset (Tabula Sapiens v2)^27^. In this setting, embeddings are produced at the resolution of individual cells: each observed expression profile is passed through GEM-1’s expression encoder and mapped into both the technical and biological latent spaces, yielding distinct representations per cell. (By contrast, for bulk RNA-seq, the embeddings represent entire samples rather than individual cells.) For scGPT^19^ (**Fig. 4A**, bottom row), embeddings were extracted by passing observed single-cell expression profiles through the encoder of the pretrained scgpt_full model, again producing cell-level representations. Embeddings were obtained using the scGPT library tasks.embed_data function from scGPT version 0.2.4. We then visualized the data similarly to the bulk RNA-seq embeddings.

**Classifier accuracy estimation.** To estimate the performance of GEM-1’s internal classifiers across each holdout dataset (**Fig. 3B**), each classifier was given latent representations derived from encoded counts to predict label probabilities for every sample in the holdout sets (e.g., predictions were based on counts alone, not metadata). The probabilities for all possible labels were ranked in descending order for each sample to determine the rank of the label in the relevant metadata. Ranks were then aggregated into four categories: rank 1, rank 2, rank 3, and below rank 3.

**Cell type annotation.** We evaluated cell type annotation performance (**Fig. 4C-D, Fig. S4**) on the holdout data (Tabula Sapiens v2), which was not observed during training of GEM-1 or any other models evaluated here. GEM-1, scGPT, and SCimilarity observed Tabula Sapiens v1 during training. (We explicitly designated Tabula Sapiens v2 as holdout data, while Tabula Sapiens v2 was released after the other models evaluated here were trained, ensuring that these constitute true holdout data for all models.) Tabula Sapiens v2 spans multiple tissues, each composed of samples belonging to one of many possible cell types, from multiple donors. The annotation task was defined as predicting the correct cell type label for each cell within its tissue context using only gene expression data (counts) as input.

We predicted cell type annotations for data from the new donors added in v2 of the Tabula Sapiens dataset using only gene expression data (counts) as input to each model as follows. For each model, we extracted cell-level embeddings using the publicly available pretrained model weights (see below for where these were obtained). A linear probe classifier was then trained for each model on its embeddings to assign cell type labels. Probe training used an 80/20 train/test split within each tissue, where the probes were trained using a portion of the holdout data (Tabula Sapiens v2). Probes were stratified by cell type whenever multiple classes were present to ensure balanced evaluation across categories. Logistic regression (implemented with scikit-learn; maximum of 1,000 iterations, random seed fixed at 99) was used as the probe. Evaluation was conducted on holdout cells not seen during probe fitting, and if a tissue contained only a single cell type in the training set, then metrics were not computed. Note that only gene expression data (counts)–no metadata–were used as input to each model (including GEM-1) for embedding generation in order to ensure a fair comparison between models.

The weights for the other pretrained models that we assessed were obtained as follows. SCimilarity^28^, using pretrained weights from Zenodo v1.1 (https://zenodo.org/records/10685499/files/model_v1.1.tar.gz); scGPT, using publicly released pretrained weights (https://drive.google.com/drive/folders/1eNdHu45uXDHOF4u0J1sYiBLZYN55yytS); and scBERT^17^, using pretrained weights from the official GitHub repository (https://github.com/TencentAILabHealthcare/scBERT). For each model, embeddings were directly extracted from the pretrained encoders and passed to a logistic regression probe that we trained for each model using the 80/20 train/test protocol described above.

Two baseline conditions were also included in the analysis. The “random” model assigned each cell a randomly generated embedding of the same dimensionality as the learned representations, providing a lower bound on classifier performance. The “baseline” approach was a GEM-1 based method in which predictions were made directly from raw counts using the pretrained cell type classifier, without a linear probe.

Performance of each model at cell annotation was assessed using macro-averaged precision, recall, and F1 score across tissues. Precision measures the proportion of cells predicted to be of a given type that were actually correct, while recall measures the proportion of cells of a given type that were successfully identified. The macro averaging scheme ensures that all tissues contribute equally to the final metric, regardless of the number of cells they contain. The macro F1 score summarizes performance by combining precision and recall into a single value that is high only when both are high and decreases sharply when one is low. This F1 score provides a balanced measure of annotation accuracy across diverse tissues and cell type compositions.

### Synthetic cohort generation and analysis

**Data processing for GTEx and TCGA.** Bulk RNA-seq data was obtained from the database of Genotypes and Phenotypes (dbGaP)^51^, specifically the dbGaP collection of the Genotype-Tissue Expression (GTEx) project^30^ and The Cancer Genome Atlas (TCGA) project^52^. These FASTQ files were then processed using the method described above for training dataset construction. Data from dbGaP were not used for model training.

**SYNTH-TEx cohort generation.** The SYNTH-TEx cohort was generated to match 30 general primary tissues in the GTEx project, with 100 samples from each sex for each tissue present in both sexes or 100 samples from the relevant sex for sex-specific tissues. These tissue and sex values were used as input metadata along with “healthy” for disease and “primary tissue” for sample type to generate expression data using GEM-1 in a fully synthetic mode (no reference conditioning).

**Comparison between SYNTH-TEx and GTEx.** For direct comparison to GTEx samples that were reported to be healthy primary tissue in dbGaP, the same number of SYTH-TEx samples were randomly selected so that tissue and sex numbers were matched across lab-generated and AI-generated samples. All analyses were performed in python 3.12 on log(CPM+1) expression data. For UMAP analysis (**Fig. 5A-B**), embeddings were calculated with the umap-learn package on GTEx samples and used to project SYNTH-TEx into the same latent space. Additional analyses and associated methods for the analyses are described here: https://app.synthesize.bio/notebooks/8ea5c433-5c2e-40ba-a2e2-48e353a5e917

**SYNTH-interferon cohort generation.** The SYNTH-interferon cohort of 100 healthy subject blood samples and 100 SLE patient blood samples was generated with metadata values comparable to the harmonized metadata for the training and out of time datasets that we compared to. Additional analyses, associated methods for the analyses, and datasets used for comparison can be found here: https://app.synthesize.bio/notebooks/8285fa3b-1647-4db8-b986-ec87828b8eec#e5abe8c7-61ae-4d01-bcdd-b71ed82e5654

**Comparison between SYNTH-interferon and clinical cohorts.** The SYNTH-interferon cohort was compared to healthy or SLE clinical samples on a per-gene basis by plotting distributions of log(CPM+1) expression data for selected interferon-induced genes (**Fig. 5C**). The cohorts were compared on a gene set basis by calculating the GSEA test statistic for each gene set in the Connectivity Map (CMAP)^53^ cytokine-induced gene expression change database for each patient cohort and plotting those scores colored by whether or not a given gene sets’ cytokine was a type-1 interferon (**Fig. 5D**).

**SYNTH-cancer cohort generation.** The SYNTH-cancer cohort was generated by parsing tissue of origin, pathological classification (cancer status and type), and subject demographics (age, sex, ethnicity) for each of the 10,523 RNA-seq-profiled samples in the TCGA Pan-Cancer Analysis Project^52^ and using these values as metadata inputs to generate expression data using GEM-1.

**Comparison between SYNTH-cancer and TCGA.** All analyses were performed in R v4.5.0. Clustering and dimension reduction analyses were performed on log(CPM+1) data. For global clustering, genes were filtered for minimum expression level (at least 10 counts in 10 samples) and then to the top 1000 most variable genes as assessed by coefficient of variation of logCPM data. Cross-substudy batch effects were removed with limma’s removeBatchEffect function^54^. UMAP dimension reduction was performed with the UMAP package using the “naive” method. Molecular subtyping was performed according to the original published method using the CMScaller R package^55^. Additional analyses and associated methods for the analyses are described here: https://app.synthesize.bio/notebooks/4a2e7d7b-cd98-473e-b088-7c6d153eafb5

### Model architecture, training, and inference

**Model architecture.** To model the relationship between tabular metadata and gene expression, we fit a deep latent variable model (DLVM) similar to a Variational Autoencoder (VAE) that employs latent generative factors parameterized by a neural network that takes the relevant metadata as input to parameterize a distribution over observed gene expression values (i.e., the decoder). We consider three independent generative factors–*biological*, *perturbational*, and *technical*–that each have independent Gaussian priors parameterized by a neural network. For example, *biological* metadata includes sex, disease, cell line, cell type, tissue, and age; *technical* includes library layout, library preparation, and platform; and *perturbational* includes perturbation embeddings, perturbation types, doses, and times.

**Model training.** The model has a likelihood Pr(expression | z_bio_, z_pert_, z_tech_) and priors Pr(z_bio_ | biological metadata), Pr(z_pert_ | perturbational metadata), and Pr(z_tech_ | technical metadata). Instead of directly optimizing the marginal likelihood, we introduce a variational posterior Q(z_bio_, z_pert_, z_tech_ | expression) (i.e., the encoder) and optimize the variational objective jointly with respect to prior, posterior, and likelihood parameters. Specifically, we maximize the evidence lower bound (ELBO) using amortized variational inference, i.e., using neural networks to parameterize Gaussian distributions over generative latent factors z = (z_bio_, z_pert_, z_tech_) as functions of the observed expression data, where our Q assumes that each generative factor is conditionally independent given expression.

**Model inference.** Notably, our modeling and inference choices give rise to multiple conditional sampling strategies that enable *in silico* generation of gene expression data that is consistent with a biologist’s expectations. In one scenario, a biologist may want to generate samples that have technical (i.e., batch) effects similar to data they currently have. Here, one samples z_tech_ from posterior Q conditioned on observed data to match the technical conditions of the observed data and then samples z_bio_ and z_pert_ from their respective priors to capture heterogeneity consistent with those factors’ metadata specifications. With realized samples for each generative factor, one proceeds with sampling the decoder for *de novo* samples that have similar technical effects as the reference data but with realistic biological and perturbational heterogeneity. In another scenario, a biologist may want to simulate what effect a perturbation might have on a set of observed data (e.g., a patient’s tissue sample). Here, one samples z_bio_ and z_tech_ from posterior Q conditioned on the observed data to match both the sample’s specific biology as well as the technical effects of the utilized sequencing technology. The biologist can then sample many z_pert_ from Pr(z_pert_ | perturbational metadata) for a wide variety of perturbational configurations and proceed with the decoder to simulate each perturbation’s effect. To the best of our knowledge, our model is the first to support such conditional sampling strategies in addition to full prior sampling (i.e., using metadata alone).

Our ability to combine posterior and prior sampling enables us to selectively control for the various types of heterogeneity: *biological*, *perturbational*, and *technical*. In the aforementioned scenarios, sampling a generative factor from its associated prior ideally captures the heterogeneity consistent with that factor’s metadata. For example, we expect reductions in prior entropy as more metadata is specified. For instance, Pr(z_bio_ | sex=M) will have higher entropy than Pr(z_bio_ | sex=M, disease=colon cancer). The former captures healthy males and diseased males across all disease ontologies, whereas the latter describes just males with colon cancer. On the other hand, the variational posterior conditioned on a reference sample will generally produce low entropy distributions that capture uncertainty over the respective generative factors for an observed reference sample.

This flexible control in using the prior for realistic heterogeneity and the posterior for adherence to a specific sample is enabled by the model’s design and is a direct result of the Evidence Lower Bound (ELBO) optimization. This ELBO decomposes locally into:

E_Q_[log Pr(expression | z_bio_, z_pert_, z_tech_)] -

D_KL_(Q(z_bio_ | expression) || Pr(z_bio_ | biological metadata)) –

D_KL_(Q(z_pert_ | expression) || Pr(z_pert_ | perturbational metadata)) –

D_KL_(Q(z_tech_ | expression) || Pr(z_tech_ | technical metadata)).

The first term’s (reconstruction) expectation is with respect to Q, the variational posterior, which is conditioned on the observed expression. The ELBO rewards large reconstruction likelihoods, which encourages Q to learn low-entropy distributions over the latent factors as a function of an observed sample. Because we fit likelihood, variational, and prior parameters jointly, the KL divergences ensure that low-entropy variational distributions will be placed in regions with high prior mass; that is, we expect all expression values with identical metadata to produce variational posteriors that fall under the Gaussian umbrella specified by the prior’s mapping of that metadata to a mean and covariance. For example, we expect Q(z_bio_ | expression) for all male samples to fall under high-mass regions of Pr(z_bio_ | sex=M). In this sense, Q(z_bio_ | expression) will produce a low-entropy distribution over the biological generative factor that promotes good reconstruction of the observed expression, akin to a VAE’s reconstruction. Here, Q’s entropy represents epistemic (posterior) uncertainty of a sample’s generative factor. In this sense, Q(z_bio_ | expression) learns a low-entropy distribution that captures the specific biological factors of an individual sample, ensuring accurate reconstruction. This low entropy reflects the model’s epistemic uncertainty about that sample. In contrast, the prior Pr(z_bio_ | sex=M) has a wider covariance that captures the full range of biological heterogeneity observed in all male samples. This complementary design gives the biologist a powerful choice: use the posterior to condition the model on a reference sample’s exact characteristics or use the prior to generate diverse, realistic samples for any metadata combination.

Another key advantage of our model’s modular latent space is its ability to transfer perturbation effects across different contexts, even when using observed data. A biologist might want to simulate the effect of a specific drug on a unique patient sample without needing to perform a physical experiment. Our model enables this by combining latent factors from multiple sources. For instance, one could take the variational posterior Q(z_pert_ | expression_drug_) derived from a reference sample treated with the drug. This captures the drug’s specific effect, including any off-target or context-dependent responses. This Q(z_pert_) can then be combined with the biological and technical posteriors Q(z_bio_ | expression_patient_) and Q(z_tech_ | expression_patient_) from the patient’s own tissue sample. By feeding this new combination of latent factors into the decoder, the model generates a new gene expression profile that represents the patient’s sample after virtual drug treatment. This recombination of posteriors allows the model to predict how a perturbation might behave in a new context, generalizing beyond the conditions seen in the original training data.

In addition to generating *de novo* gene expression, we are able to predict metadata relevant to each generative factor from the factors themselves. We found that adding linear classifiers and their resulting likelihoods to the ELBO encourages a latent space that is linearly separable according to metadata, which we find greatly improves model performance. Additionally, this ability enables biologists using our model to impute missing and/or detect mislabeled metadata for a set of observed expression data.

**Perturbation representation and embedding extraction.** The selection of diverse molecular embedding models is crucial for enabling our model to generalize effectively to new perturbations. The choice of embeddings was driven by the principle of capturing a comprehensive and complementary set of molecular features, and the specific combination was selected by hyperparameter optimization.

To model the effects of perturbations on gene expression, we utilized structured metadata annotations for each sample. The perturbation ontology identifier field indicated whether a sample was perturbed and specified the perturbing agent, while perturbation type denoted the perturbation mechanism: small molecule exposure or gene-level modulation via shRNA knockdown, CRISPR knockout, or gene overexpression. Perturbation identifiers were normalized using standard ontologies. Genes were mapped to Ensembl genes (ENSG identifiers), small molecules to ChEBI, and chemical compounds to ChEMBL synonyms. For each perturbation, we extracted pretrained embeddings based on its type.

For **molecular perturbations**, we used embeddings from the following foundation models:

- **ConPLex**^56^ – contrastive protein-ligand representation model
- **RDKit2DNormalized** – 2D structural descriptors generated using RDKit
- **ErG**^57^ – Extended reduced graph fingerprints
- **PubChem fingerprints** – binary structural fingerprints from the PubChem database
- **Morgan Fingerprints**^58^ – circular topological fingerprints
- **MoleculeSTM**^59^ – transformer-based self-supervised molecular embeddings

For **gene-based perturbations**, we leveraged precomputed embeddings^60^ from the following approaches:

- **MUT2VEC**^61^ – unsupervised gene embeddings from mutational co-occurrence
- **BIOCONCEPTVEC-CBOW**^62^ – biomedical concept-based word embeddings
- **ESM2**^3^ – transformer-based protein language model trained on UniRef50
- **Node2Vec**^63^ – graph embeddings derived from protein-protein interaction networks

In instances where a perturbation could not be mapped to any pretrained embedding (e.g., due to missing ontology alignment), we applied one-hot encoding to represent the perturbation identifier. This fallback ensured that all samples had a defined perturbation representation, allowing the model to learn from the full dataset.

## Data availability

All bulk RNA-seq datasets used for model training are publicly available through the NCBI Sequence Read Archive (SRA)^12^. We obtained protected GTEx^30^ and TCGA^52^ data from dbGaP^51^ and used these datasets for evaluation purposes only, not for model training. Single-cell RNA-seq datasets used for model training and evaluation are publicly available from CZI CELLxGENE Discover^16^ and the scPerturb database^25^ of harmonized SRA and Gene Expression Omnibus (GEO)^64^ data. The AI-generated synthetic cohorts described here (**Fig. 5**) are freely available for download and re-use (https://app.synthesize.bio/datasets).

## Model and code availability

The model code and weights will be made available in a future version of this manuscript.

## Acknowledgements

The results published here are based in part on data generated by the TCGA Research Network: http://cancergenome.nih.gov/. TCGA data used for analyses described in this manuscript were obtained from dbGaP accession number phs000178.v11.p8 on 11/30/2024. The Genotype-Tissue Expression (GTEx) Project was supported by the Common Fund of the Office of the Director of the National Institutes of Health, and by NCI, NHGRI, NHLBI, NIDA, NIMH, and NINDS. GTEx data used for the analyses described in this manuscript were obtained from dbGaP accession number phs000424.v10.p2 on 11/30/2024.

## Author contributions

Using the CRediT system:

Conceptualization: GK, AMW, JTL, RKB

Data curation: AMW, KAJ, CS

Formal analysis: VM, KAJ, MH, ARA, S-YN

Investigation: GK, S-YN, VM, KAJ, MH, ARA

Methodology: GK, AS

Project administration: GK, AMW, LJR, SS, JTL, RKB

Resources: AD, DH, JMK, LJR, SS

Software: AD, DH, JMK, SS

Supervision: GK, AMW, JTL, RKB

Validation: VM, MH, KAJ

Visualization: VM, MH, KAJ, ARA

Writing – original draft: GK, AMW, VM, KAJ, MH, ARA, JTL, RKB

Writing – review and editing: GK, AMW, VM, KAJ, MH, ARA, ARB, AD, DH, JMK, S-YN, LJR, CS, SS, JTL, RKB

---

**Figure S1.**
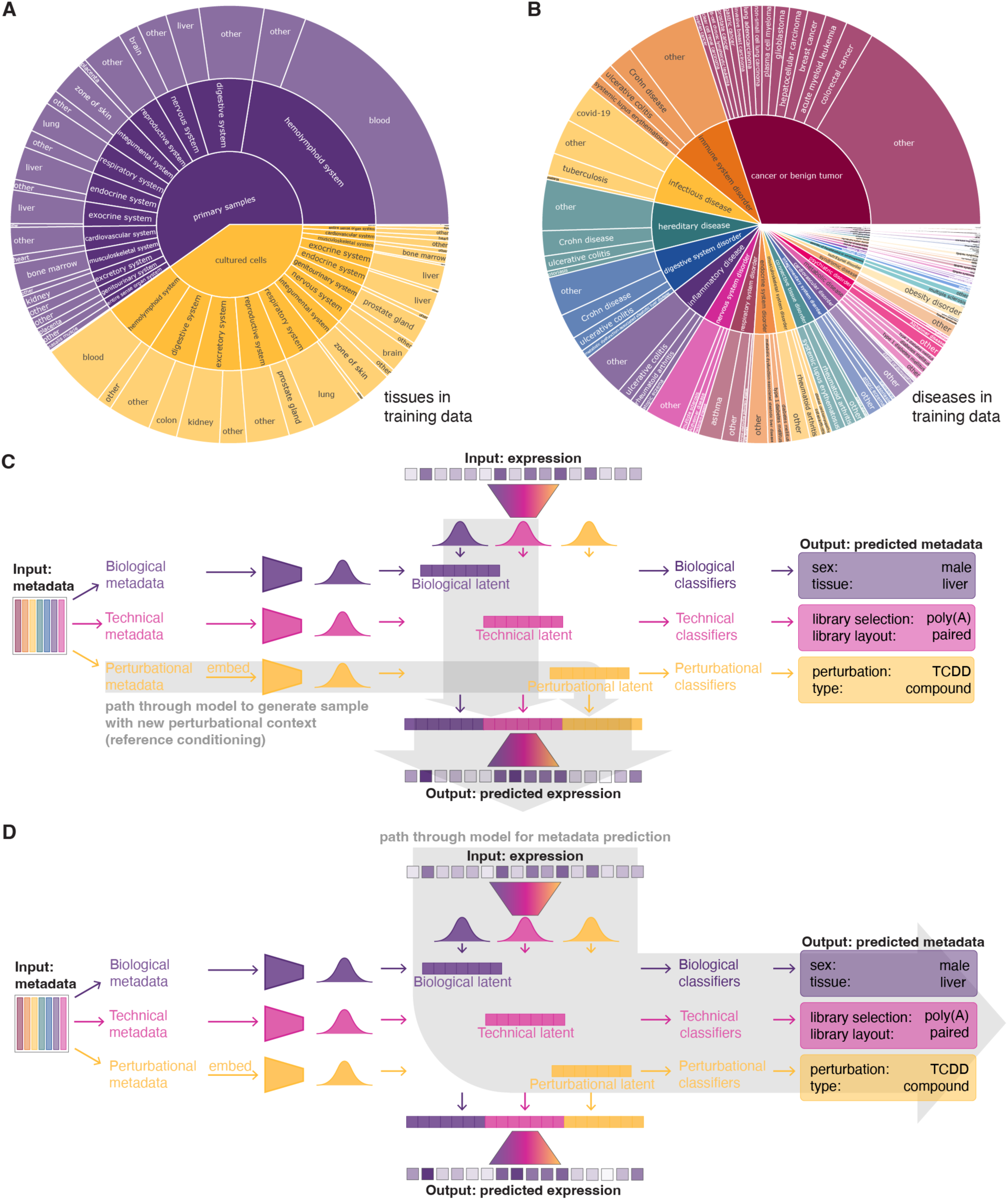
-related to Figure 1. Distribution of training data and model invocations for different inference problems. (**A**) Sunburst plot illustrating the distribution of bulk RNA-seq training data among tissue types for cell lines (orange) and primary samples (purple). Other sample types (e.g., organoids) are not included for clarity of illustration. Note that a given sample can belong to multiple categories. (**B**) As (A), but for diseases. (**C**) Schematic illustrating GEM-1’s architecture and path through the model for generating a sample with a new perturbational context (e.g., generating a perturbed sample conditioned on expression data for an unperturbed, control sample). (**D**) As (**C**), but illustrating the path through the model for predicting metadata conditioned on expression data.

**Figure S2.**
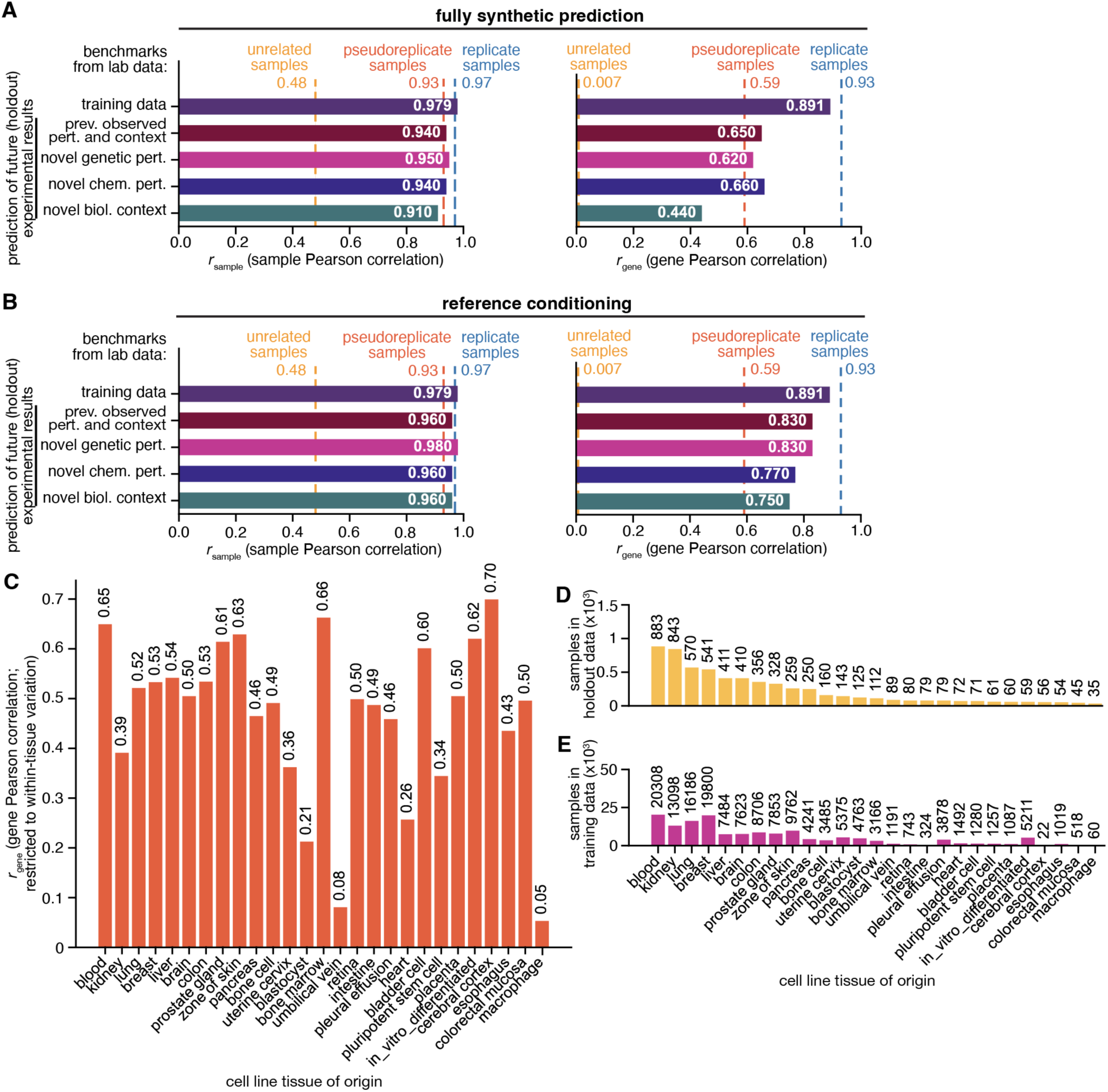
-related to Figure 2 Evaluation of model performance across different experimental settings and biological contexts. (**A**) As Fig. 2E, but additionally illustrating *r*sample (sample Pearson correlation). (**B**) As Fig. 2F, but additionally illustrating *r*sample (sample Pearson correlation). (**C)** Bar plot illustrating *r*gene for fully synthetic prediction of holdout data, computed by stratifying cell lines based on their tissue of origin and then calculating *r*gene across those cell lines. Because this computation is performed on a within-tissue basis, the resulting *r*gene values reflect only within-tissue variation and not cross-tissue variation, which is typically the primary source of biological variability in gene expression. (This is why the values plotted here are lower than those in (A), which includes cross-tissue variation in the computation.) (**D**) Bar plot illustrating the numbers of samples in holdout data for each tissue type analyzed in (**C**). (**E**) As (**D**), but for training data.

**Figure S3.**
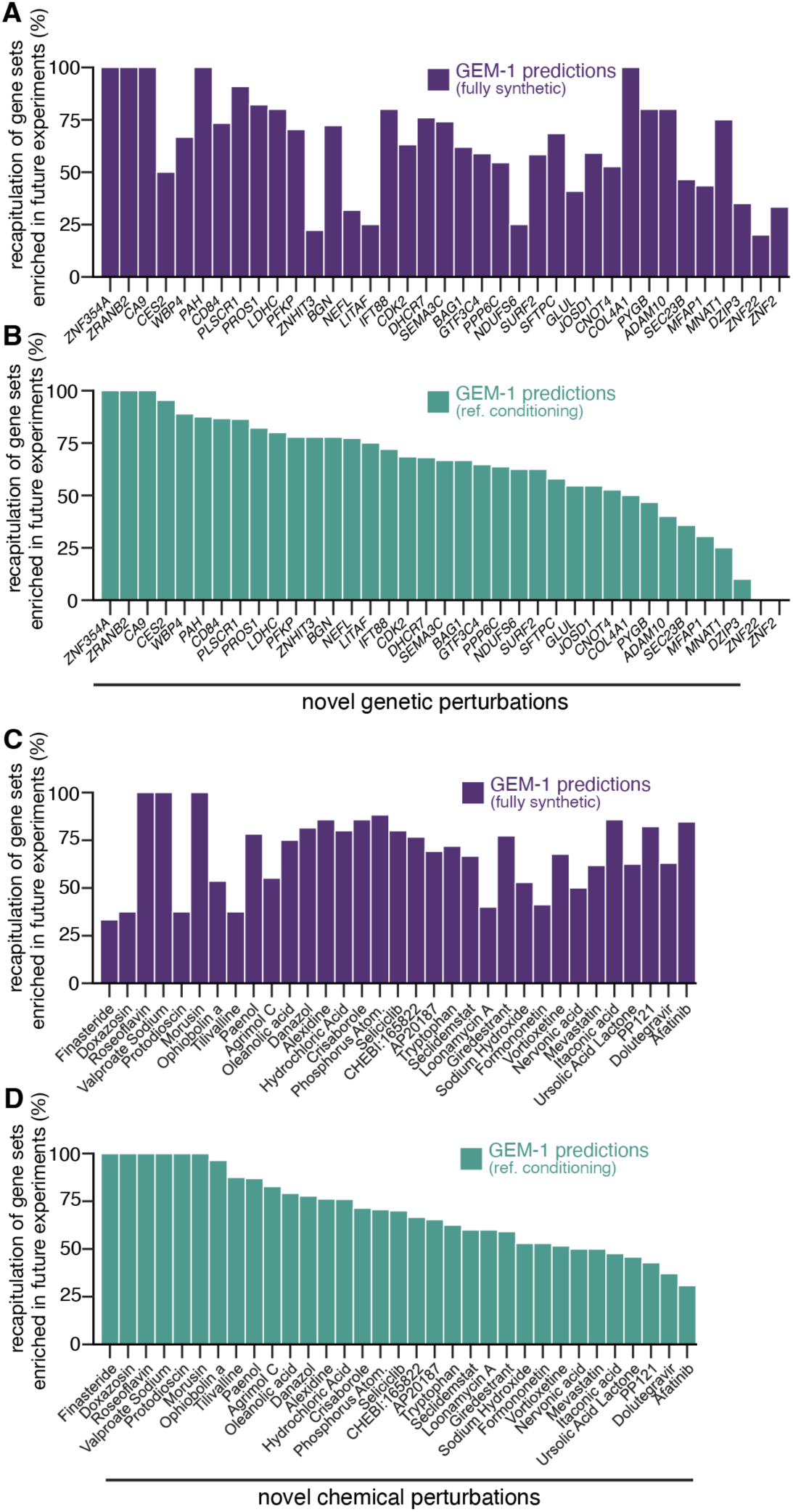
-related to Figure 2. Evaluation of GEM-1’s ability to recapitulate key molecular phenotypes following genetic or chemical perturbation. (**A**) As Fig. 2J, but for fully synthetic AI predictions. (**B**) Identical to Fig. 2J; reproduced here for comparison purposes. (**C**) As Fig. 2J, but for fully synthetic AI predictions made for novel chemical perturbations. Plot restricted to the 33 chemical perturbations for which we identified metadata-matched control (unperturbed) and perturbed samples in holdout data. (**D**) As (**C**), but for AI predictions with reference conditioning on control (unperturbed) samples.

**Figure S4.**
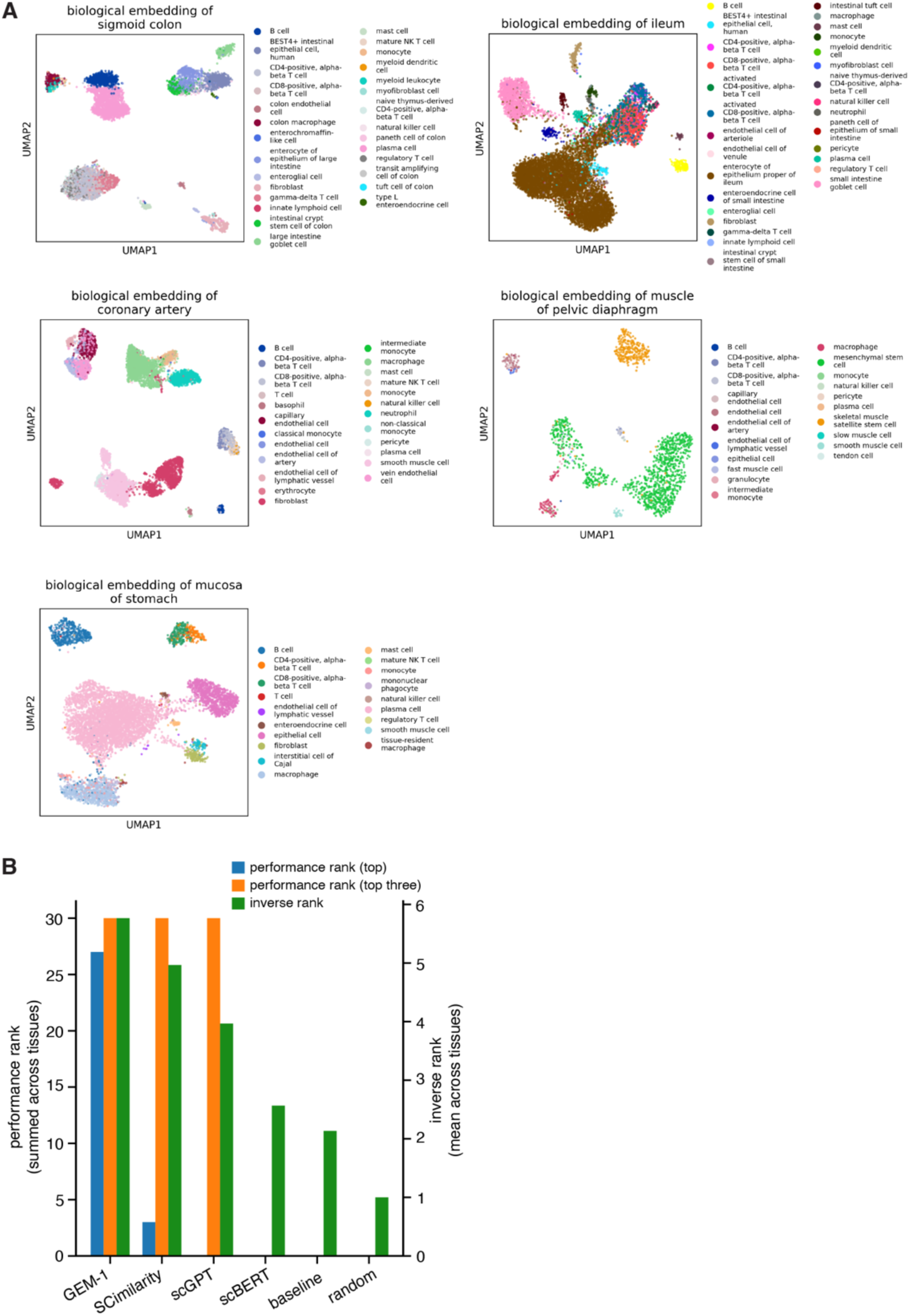
-related to Figure 4. GEM-1 effectively annotates cell type in single-cell data. (**A**) As Fig. 4A, but with legends indicating the mapping from colors to cell types. (**B**) Bar plot illustrating the performance of different models when used for cell type annotation. Plot is derived from the macro F1 analysis illustrated in Fig. 4C.

